# Delaying the onset of aided target recognition highlights allows for a more dispersed allocation of overt attention

**DOI:** 10.64898/2026.06.30.735590

**Authors:** Chloe Callahan-Flintoft, Angela Jeter, Jessica L. Villarreal, Gabriella B. Larkin

## Abstract

Visual search is a critical component of many professions such as military operations, baggage screening, and radiology. Aided Target Recognition (AiTR) systems are designed to highlight potential threats across the operator’s visual field in real-time, directing attention and improving accuracy. However, these systems may impact search and, consequently, situational awareness by diverting attentional resources from non-highlighted, yet relevant, locations. Previous work suggests that scene gist is extracted within the first 250 ms of scene onset (Vö & Henderson, 2010). As such, this study examined whether a 250 ms AiTR onset delay could encourage a more even distribution of attention. Participants searched synthetically generated scenes and classified each person in the scene as armed or unarmed. Depending on their condition, participants either saw the scenes unaugmented (No AiTR condition), with AiTR highlights consisting of red bounding boxes around armed people and yellow boxes around unarmed (AiTR condition), or with AiTR highlights presented 250 ms post scene onset (Delayed AiTR condition). A surprise memory test of background objects presented in the search scenes was administered to all participants upon completion of the search task. As predicted and preregistered, results showed less overt attentional deployment to background information (anything other than the people themselves) in the AiTR condition compared to No AiTR, however, decreased overt attentional deployment was not seen in the Delayed AiTR group. A similar pattern was observed in the memory data (with the AiTR condition having a lower score than the No AiTR condition and the Delayed AiTR condition), this difference was not significant.

**Significance statement:** Aided target recognition (AiTR) systems are designed to augment a user’s field of view with salient highlights at locations containing task-relevant or threatening information in the environment. As computer vision models advance, such technologies will become an even more critical tool for people that perform visual search professionally (e.g. Soldiers, baggage screeners, radiologists). In a military context especially, having a system that potentially improves threat detection accuracy and reduces response times could be lifesaving. However, such benefits can only be realized if AiTR systems are designed to be integrated with the cognitive mechanisms of the human user. Without this, displays run the risk of being distracting, creating attentional blind spots, and generally producing a poorer user experience and performance. The current work, motivated by findings in the literature, tested the introduction of a short delay between scene onset and the onset of AiTR highlights. It was found that this very brief delay (250 ms) was sufficient to allow participants to direct overt attention to areas of the scene not highlighted by the AiTR system. This finding is critical because, while the purpose of AiTR highlights is to encourage the placement of attentional resources across the visual field, it is imperative that the human operator ultimately be able to attend to non-highlighted locations in the event of system failures (e.g. threats missed by the AiTR). Moreover, this work demonstrates how cognitive research can be used to systematically explore complex application spaces for improve human-machine integration.

## Introduction

There are a variety of professions in which visual search is a key component and where failure to detect a target could have profound consequences (e.g. soldiers monitoring a checkpoint, baggage screeners inspecting bags for prohibited items, radiologist scanning CT scans for abnormalities). As such, there is an ongoing effort to develop technological solutions to aid in these critical search tasks. In a military context, these systems are referred to as Aided Target Recognition (AiTR). With mobile AiTR systems, a computer vision algorithm, via some heads-up display (e.g. augmented reality glasses, within a rifle scope, etc), displays salient highlights on locations in the human operators’ visual field where objects of interest (e.g. potential targets or threats) are likely present. As with other attentional cueing technologies, such as Computer Aided Detection (CAD) systems used in the field of radiology (Castellino, 2005), the goal is to increase detection rate and accuracy. Specifically in a military context, another key performance metric for AiTR is to draw the operator’s attention to locations of interest faster than would occur naturally. For soldiers, who are often doing multiple tasks simultaneously (e.g. monitoring for threats while also navigating a mission objective), an AiTR system’s assistance with threat detection could free up cognitive resources tied to target search, allowing them to be reallocated to other visual cognitive tasks. However, AiTR displays must be developed within the context of human cognitive mechanisms as failure to properly integrate such displays the processes and timescales of visual perception will impede the ability to achieve the desired effects. These challenges are similarly found in the field of radiology where CAD systems, intended to assist radiologists detecting the presence of cancer nodules in CT scans, result in very little net gain for detection and identification (Fenton et al., 2011; Fenton et al., 2007). Specifically, miss rates increase for targets that CAD systems fail to highlight (Drew, Cunningham, & Wolfe, 2012). This decreased ability to detect unhighlighted targets is of particular importance in a military context as reduced attention to background/unhighlighted stimuli can have disastrous implications for situational understanding in life and death circumstances. Additionally, new operational environments may degrade algorithm robustness and evolving adversarial actions may change the accuracy, both correct detection and false alarms, of any future AiTR system. As such, systems must be designed with the assumption that they will occasionally fail (e.g. miss a target) and the human operator will need the ability to maintain performance levels in that event. Such systems therefore must be designed to increase rate and speed of detection, all without compromising the soldier’s situational awareness or impeding the soldier’s ability to detect targets missed by the system. To do this, AiTR system design must exploit and support the way in which the human visual system processes information.

Understanding how AiTR highlights affect visual search first requires an understanding of how search is performed in an un-augmented context. Real world scenes contain a complex array of visual information that cannot be instantaneously processed at a uniform level of accuracy. To build meaningful internal representations of the world around us while minimizing processing lag, the visual system must prioritize some information for enhanced processing. This is done through a combination of pre-attentive processes, covert attentional shifts, as well as overt eye movements. While covert and overt attentional systems are not strictly coupled in their deployment of resources (Talcott & Gaspelin, 2021), they do show similarities in what factors influence their deployment. Many models or conceptualizations of attention divide these factors into three categories: bottom-up saliency signals, top-down control settings implemented by the participant based on the task goals at hand, and selection history effects (Theewues, 2019). Salient items in the visual field are known to capture attention, whether by slowing response times to a non-salient target (Bacon & Egeth, 1994; Theeuwes, 1992) or even perturbing saccade trajectories (Godijn & Theeuwes, 2002). Moreover, research has found that, relative to scene onset, early eye movements are often guided by these saliency signals (Anderson et al., 2015). However, when using real world scenes as stimuli, while saliency models account for a significant portion of the variance in participants’ fixation patterns, they fail to provide complete predictions of where people fixate (Foulsham & Underwood, 2008).

As is evident by participant’s ability to ultimately find the target, even in the presence of a salient distractor (e.g. Lamy, Tsal, & Egeth, 2003), eye movements are not solely driven by bottom-up signals. In fact, gaze pattern significantly alters with instructional set (Siebold & Donk, 2014; Yarbus, 1967). When bottom-up and top-down influences conflict, there are a variety of mitigating factors that can determine where the eyes move. For instance, the eyes are more susceptible to move towards salient distractors when participants have weaker top-down control (e.g. when they are tasked with finding the odd shape in the array, as in a pop-out search, versus when they know what the target looks like, as in a feature search) (Gaspelin, Leonard, & Luck, 2017; Gaspelin & Luck, 2018). Moreover, evidence suggests that when cognitive resources are strained, potentially affecting one’s ability to implement top-down control, the brain uses saliency to prioritize information for encoding (Fine & Minnery, 2009), which is consistent with findings that individuals with lower working memory capacity are more prone to attentional capture by salient distractors (Gaspar et al., 2016).

There is an important component when studying vision of real-world scenes that is missed with the more tightly controlled, abstract displays used in many experimental paradigms. Namely, real-world scenes, in and of themselves convey meaningful information, even in the absence of a specific task or goal. As such, meaningful locations in a scene, or those locations that participants judge to have semantic relevancy to the scene, are highly fixated (Henderson & Hayes, 2017; Henderson & Hayes, 2018). This automatic extraction of scene gist has important implications for visual search behavior in real world images that is absent in more abstract displays, as participants are shown to use scene knowledge to guide attentional allocation in the search for targets (Krzys et al., 2023): participants were more likely to fixate on locations where a target was expected when that target had a high certainty location (e.g. a sink has a high certainty of being located in the countertop of a kitchen). This underlying structure of natural images, termed ‘scene grammar’ (Vö, Boettcher, & Draschkow, 2019) could potentially be grouped under the umbrella of ‘selection history’ effects (Theeuwes, 2019) as it is at least partially learned through experience.

Taken together, this wealth of knowledge on the factors that guide attentional allocation already provide keen insight into how AiTR highlights might affect search behavior. For one thing, AiTR highlights, by design, are highly salient. The current imagining of AiTR highlights, and implementation in similar CAD systems (Dromain et al., 2013), is a hard bounding box, usually with a learned color palette to indicate classification (e.g. enemy or friendly) or color gradient associated with algorithm certainty (e.g. red for highly certain threat down to yellow for uncertain threat). Highlights will naturally become part of a soldier’s visual search task-set as their presence should coincide with the location of objects relevant to the soldier’s task. This implies that both the bottom-up saliency of the highlights as well as their top-down, task relevancy will drive attention to highlighted locations in the visual field. The current work presented participants with simulated scenes containing people and background objects. Using mouse clicks, participants were tasked with classifying each person in the scene as having a weapon or not. Depending on the participant’s assigned condition, the scenes were either presented alone or with AiTR highlights (in the form of red and yellow bounding boxes) around each person in the scene. Given the combination in up-weighting of AiTR highlights, through both bottom-up saliency and top-down task relevancy, the first hypothesis of this experiment was that there would be less attention allocated to background objects (anything other than the highlighted locations) when AiTR highlights were present, compared to when no highlights were present. Here, attentional allocation was operationalized as the rate at which fixations were made on background information across AiTR conditions.

The point of AiTR highlights is to direct attentional resources to locations of interest in the environment. However, if attentional resources are finite, the direction of such resources by AiTR highlights necessarily comes at a cost to unhighlighted locations. As discussed earlier, no system will be perfect in detecting objects of interest. Moreover, the human operator has a unique ability to understand scene context and flexibility to re-prioritize task goals in dynamic environments. As these abilities are typically absent in computer vision algorithms, it is essential that while AiTR displays should encourage attentional allocation, these displays should not override the human’s ability to attend to unhighlighted locations. In military contexts, this more wholistic processing of a scene is often referred to as situational awareness. Therefore, the second hypothesis of this experiment is that, as a consequence of redirected attentional resources, participants will have poorer situational awareness when AiTR highlights are present compared to when they are not. To operationalize situational awareness, a surprise memory test was given to participants to assess their memory for the background items of the scenes they had previously searched.

Finally, previous work suggests that a scene’s “gist” can be extracted rapidly, within the first 250 ms of scene exposure (Vö & Henderson, 2010), with different time courses (80 ms to 500 ms) associated with different components and capabilities (Mudrik, Faivre, & Koch, 2014). Moreover, temporal models of attentional competition predict that less salient objects in the visual field can still “win” the locus of attention if they are presented before more salient objects (Wyble et al., 2020). Given these past findings and model predictions, a delayed AiTR condition was included where participants were first shown the scenes and then, 250 ms post scene onset, AiTR highlights were presented. The third, and final, hypothesis then is that delaying the presentation of AiTR highlights will allow more attentional allocation to background objects and, consequently, better memory for those objects compared to the performance of participants who were presented AiTR highlights simultaneously with scene onset.

## Methods

### Ethics statement

This experiment was approved by the Institutional Review Board of the U.S. Army Research Laboratory (ARL) under Project Number ARL 21-054. All procedures were in accordance with the Declaration of Helsinki.

### Participants

Fifty-six military veterans or active-duty members were recruited for this study (19 female with an average age of 41). All participants were recruited from the Joint Base San Antonio (JBSA), Ft. Sam Houston catchment areas and had normal or corrected to normal vision with no reports of color blindness. Participants reviewed and signed an Institutional Review Board (IRB) approved consent form prior to the experiment. Pre- and post-screen questionnaires were administered consisting of demographics, handedness, gaming experience, and strategies used. This data is not reported in the current paper however is available (along with this study’s preregistered analysis plan, experimental data and paradigm code at www.osf.io/3rka8). Eight participants were removed from analysis as over half of their eye tracking samples were invalid due to them moving their head out of range and another five participants were removed due to a technical issue whereby the marker stream, which links the eye tracking data to the experiment file, was not saved.

Participants were pseudorandomized into one of three conditions (No-AiTR, AiTR, Delayed-AiTR). The pseudorandomization was constructed such that the condition group for each participant was sampled without replacement from the possible three conditions (e.g. the first participant had equal chance of being placed in any of the three conditions, the second participant scheduled had equal chance in being placed in the remaining two condition and so forth). This was done to maintain comparable sample sizes across conditions throughout data collection. At the start of data collection a fourth condition was included, however this condition was dropped after only collecting four participants to focus recruitment efforts and maximize sample size for the main three conditions. Therefore, the results for the fourth condition are not reported or discussed here. After exclusions, 14 participants were analyzed in the AiTR condition, 15 in the No AiTR condition and 14 in the Delayed AiTR condition.

### Apparatus

Stimuli were developed using Unity (Unity, Technology Inc.) and were displayed on the Tobii Pro Spectrum monitor (1920 x 1080 pixels). The Tobii Pro Spectrum tracks eye movements using infrared emitters with a 300 Hz sampling rate. Data collected included gaze position and pupil diameter. To calibrate, participants followed instructions of Tobii’s 5-point calibration procedure.

### Procedure and Stimuli

Participants were given a COVID screening to ensure safety and all proper disinfectant precautions were administered. Participants were then assessed with the Snellen chart and a 14 plate Ishihara colorblind test prior to starting the experiment. The participants were instructed to get into a comfortable position and limit their head movement as much as possible. After eye tracking calibration was completed, participants were asked to maintain this position throughout the experiment.

For the visual search task, participants were presented with scenes containing people in four possible settings (office, train station, boardwalk, and forest) (Figure 1). Each scene contained six people (two to five carrying weapons). Five pseudo-randomly selected task-irrelevant items (e.g. pumpkin, hand sanitizer bottle, trophy) and two randomly selected task-relevant items (e.g. weapons, not carried by a person) were scattered about each scene. There were two groups of 25 task-irrelevant objects. Each participant was randomly assigned one group to be used in the visual search scenes and the other group was used as lure items in the subsequent memory test. As there were 5 objects used in each of the 100 scenes shown to participants, selected from a list of 25 objects, objects were balanced such that each appeared 20 times, never in the same scene twice. For the task-relevant items, two weapons were selected out of a total list of 10 possible weapons for each scene. These were the same 10 possible weapons that a person in the scene could be holding. Task-relevant items were selected at random for each scene and not strictly counterbalanced. As such these items appeared 16 to 22 times in the search task. The same task-relevant item never appeared twice in the same scene.

**Figure 1:**
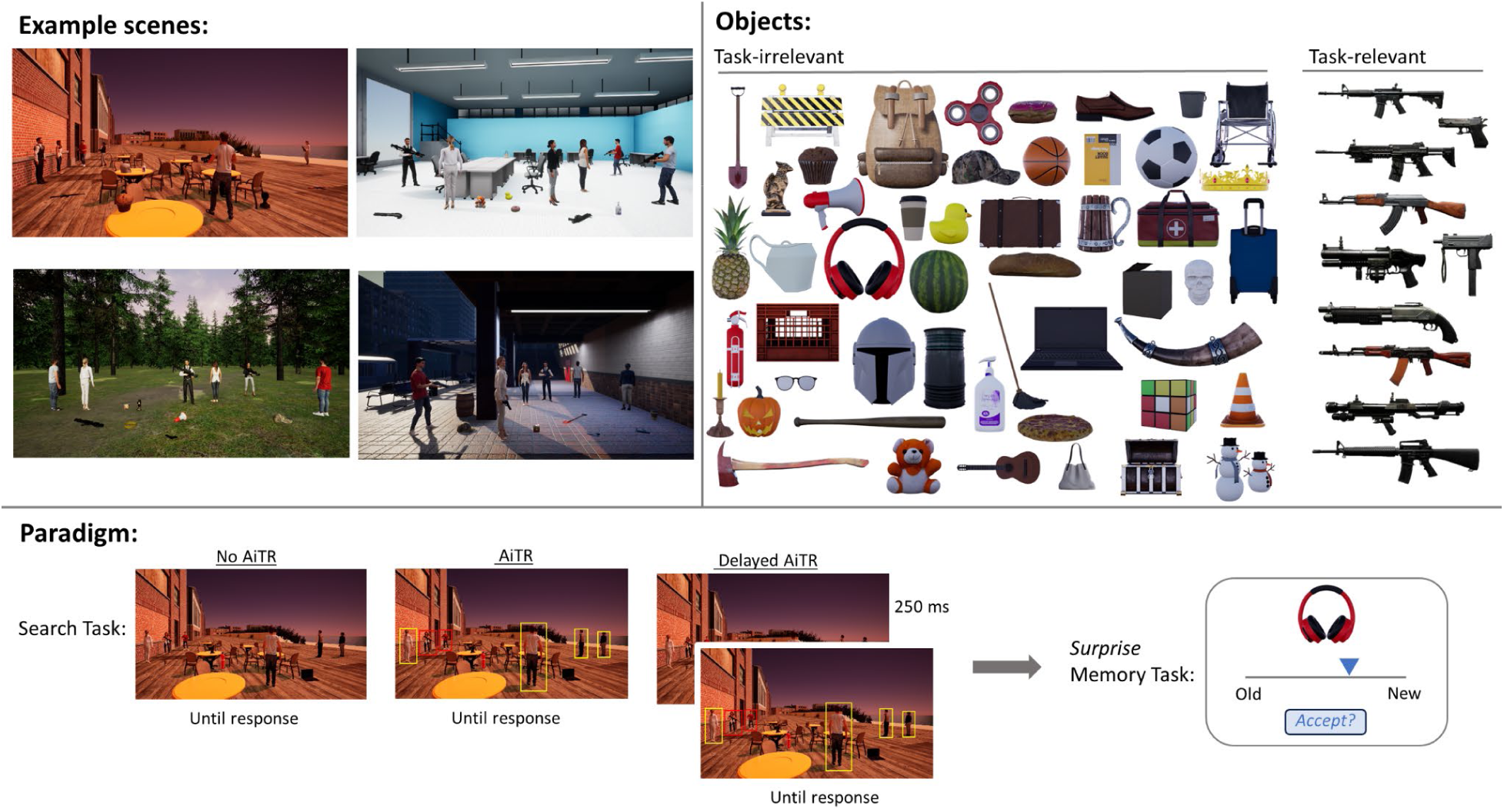
A breakdown of experimental stimuli and paradigm schematic. Top left: four example scenes, one of each background type (boardwalk, office, forest, train station). The same scenes were used across condition groups with AiTR highlights presence and presentation dependent on condition. Top right: All 50 task-irrelevant and task-relevant items used as background objects for scene construction. Participants only ever saw 25 objects in the visual search task, with the remaining 25 objects used as lures in the memory task. Bottom: a schematic of the search task followed by the surprise memory task. Participants either search the scenes alone (No AiTR condition) or with AiTR highlights presented simultaneously with scene onset (AiTR condition) or delayed in onset by 250 ms (Delayed AiTR condition). Participants in all conditions then completed the memory task where they were asked about the 25 task-irrelevant objects they saw as well as 25 lures.

Participants were instructed to find the people in each scene and to identify them as a threat (holding a weapon) by a right mouse click at their location or as a non-threat (not holding a weapon) by a left mouse click at their location, as quickly and accurately as possible. Participants in the AiTR and Delayed-AiTR conditions were told that an AiTR system, with 85% accuracy would aid them by highlighting people with weapons via a red bounding box and people without weapons via a yellow bounding box. The AiTR highlights were pre-programmed into the scene such that, throughout all of the scenes viewed, 85% of people highlighted were correctly classified and 15% were misclassified. No people were missed by the dummy-AiTR system.

The start of each trial began with a central fixation cross on a gray screen where participants were instructed to press the spacebar to begin the trial. In the No-AiTR condition, participants were presented with the unaugmented scenes. In the AiTR condition, participants were shown the same scenes augmented with AiTR highlights. Lastly, in the Delayed-AiTR condition, participants were first presented with the unaugmented scenes and then 250 ms post scene onsets, AiTR highlights appeared. In all conditions, participants classified each person in the scene as quickly and accurately as possible and hit the spacebar when they were finished, whereupon the scene would offset and they would again be presented with a central fixation cross and the prompt to begin the next trial. After the first four trials, the participant was asked if they understood the task or if they had any questions for the experimenter. These four trials were excluded from analysis as practice trials. A hundred trials were completed in total (25 with each of the four backgrounds, randomly mixed within block).

At the end of the 100 search trials participants were given a surprise memory test where they were shown each of the 50 items (25 used in the search experiment and 25 lures that were not seen previously) one by one and asked to rate on a continuous scale how “new” or “old” they felt the item was. On each trial, an item was presented with a line below. The left endpoint of the line was labeled ‘old’ and the right endpoint was labeled ‘new’. Participants were instructed to click the position of the line that best reflected their memory representation. A blue arrow would appear at the location on the scale which the participant clicked along with a button below that said ‘accept?’. Participants had the opportunity to adjust the arrow left or right on the line and click the ‘accept?’ button when they were satisfied with the rating. Old and new objects were presented in a randomly shuffled order. After all 50 objects had been rated the experiment was over.

### Analysis

Gaze position was collected in the form of x,y coordinates on the screen as well as pupil diameter. A blink detection algorithm was first run on the data detecting drop-out periods defined as samples in which the pupil diameter was reported as less than zero. If the blink or drop-out period was under 700 ms, a linear interpolation was done. If the period extended longer than that, those samples, along with the neighboring 2 samples on either side of the drop-out were marked as unusable and not included in saccade classification. Saccades were classified using the Engbert & Mergenthaler (2006) method whereby an elliptical velocity threshold is set in x,y, coordinate space per participant. This is done by taking the participant’s average velocity and then setting a velocity threshold 3 standard deviations from that average in the horizontal and vertical direction. Saccades were rejected if they were under 20 ms in duration. Fixations were defined as the time windows in between classified saccades. To test the distribution of attentional allocation, the percentage of fixations on background information was analyzed. Specifically, background fixations were denoted as fixations on anything in the scene other than the people themselves, as defined as the area within the AiTR bounding box. To equate the three conditions, only fixations made 250 ms post scene onset were analyzed, however results did not differ when this exclusion criteria was removed.

## Results

### Accuracy and response time

As accuracy was near-ceiling in this task, a non-parametric Kruskal-Wallis test was run to compare across groups. A significant difference was found between groups *χχ*^2^(2, N = 51) = 8.32, *p* = .01. This was driven by a significant difference in the mean ranks of the No AiTR condition (*M* = 0.97, *SE* = 0.003) and the Delayed AiTR condition (*M* = 0.90, *SE* = 0.04). The AiTR condition (*M* = 0.86, *SE* = 0.07) did not have a mean rank significantly different from either group. A one-way ANOVA on reaction times showed no significant difference across condition, *F*(2,48) = 0.56, *p* = 0.57.

### Attentional allocation

#### Preregistered analyses

An omnibus one-way ANOVA was run on the percentage of background fixations across conditions for completeness, *F*(2,40) = 3.38, *p* = .04, *η^2^* = 0.14. However, the specific preregistered comparisons and hypotheses were 1) a decrease in background fixations between the No AiTR and AiTR condition and 2) an increase in background fixations between the AiTR condition and the Delayed AiTR condition. A one-tailed independent sample t-test showed that there was in fact a decrease in the percentage of background fixations between the No AiTR condition (*M* = 0.32, *SE* = 0.04) and the AiTR condition (*M* = 0.21, *SE* = 0.03), *t*(42) = 2.39, *p* = 0.01, Cohen’s *d* = 0.89. Also as predicted, there was a significant increase in the percentage of background fixations between the AiTR condition and the Delayed AiTR condition (*M* = 0.31, *SE* = 0.04), *t*(42)= - 2.87, *p* = 0.02, Cohen’s *d* = -0.81 (Figure 2).

**Figure 2:**
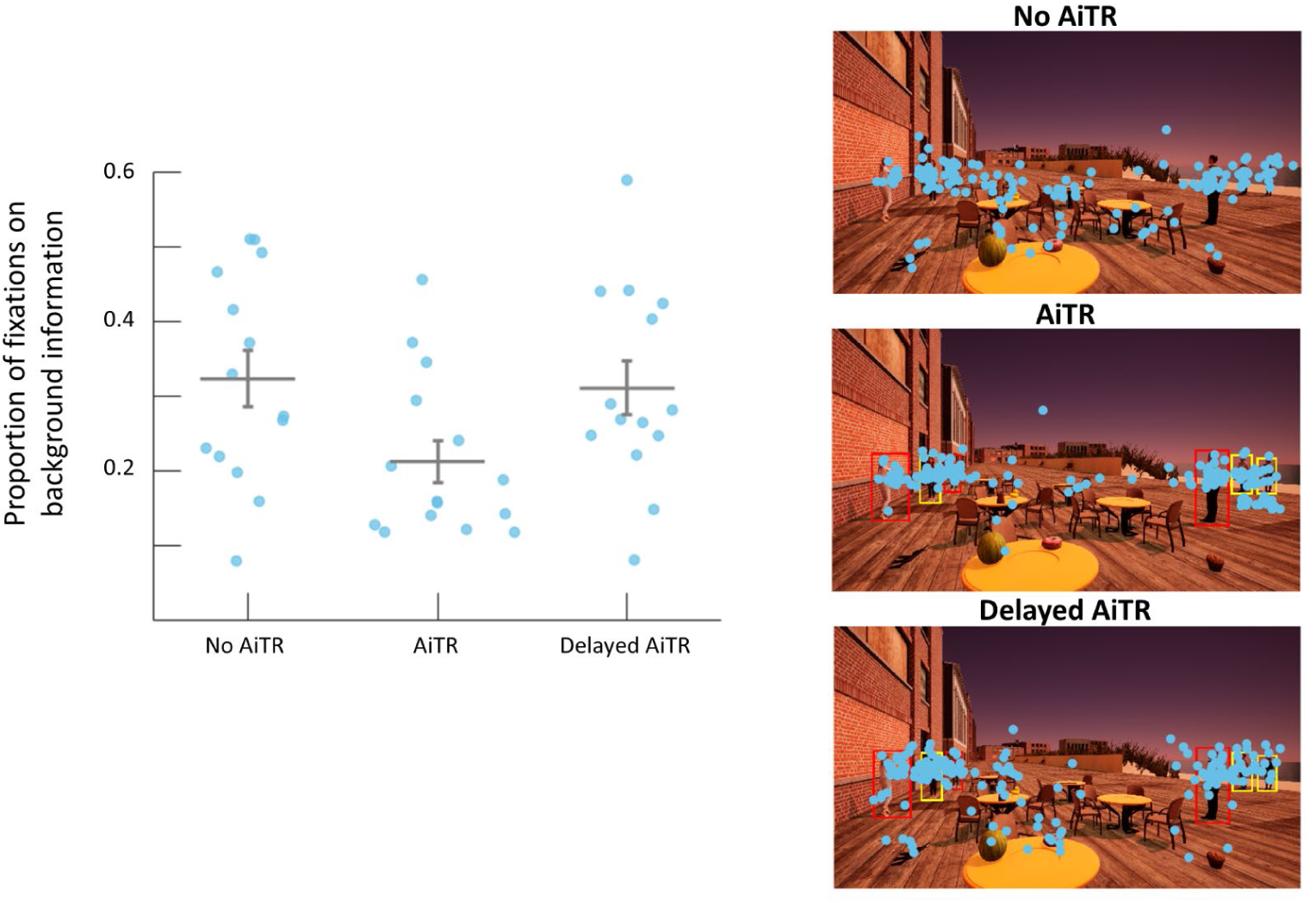
Percentage of fixations made on background information (e.g. not on the people in the scene) across conditions (left) and fixation positions overlaid as blue dots for an example scene across condition (right).

**Figure 3:**
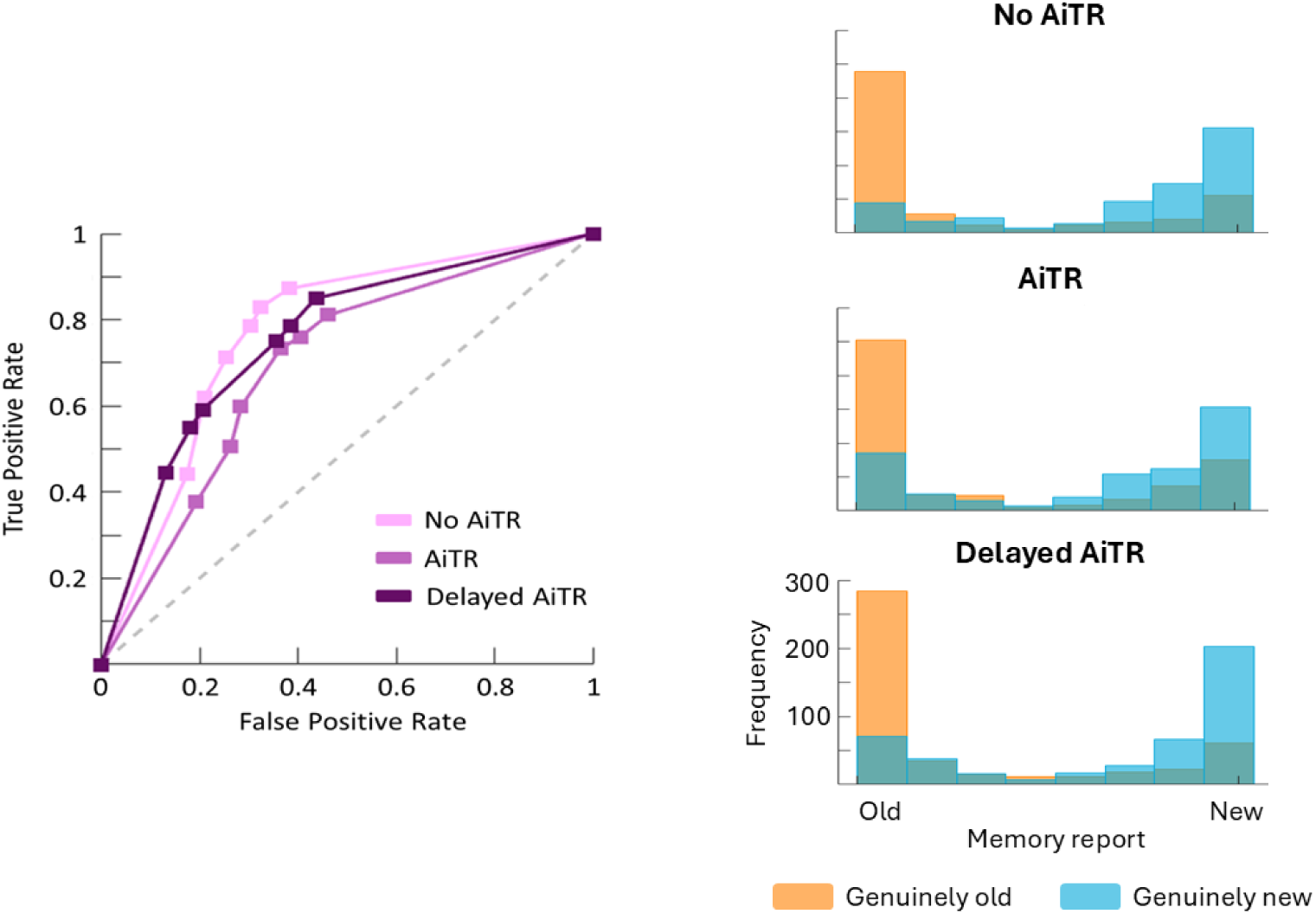
ROC curves plotted for each condition (left) and histograms of memory reports for old and new objects across conditions (right).

#### Exploratory analyses

A set of exploratory analyses were run in order to inform future research efforts. These results should be replicated in new datasets before being considered conclusive. The first of these analyses was an alternative approach for measuring the allocation of overt attention, namely, investigating whether the amount of scene coverage by fixations differed across conditions. This method, adopted from Castelhano & Heaven (2010), counts every pixel that falls within a 2 degree radius of each fixation, without counting redundant coverage, as a metric of total scene coverage. The current data set showed a similar pattern in scene coverage as was seen in the preregistered analysis, where there was less overall area covered by background fixations in the AiTR condition (*M* = .02, *SE* = 0.003) compared to the No AiTR (*M* = .03, *SE* = 0.004) and Delayed AiTR condition (*M* = .03, *SE* = 0.005). However this difference was not significant, *F*(2,40) = 2.189, *p* = 0.12.

Saccade amplitude and fixation duration on targets were also compared across conditions. While there was no significant difference across condition on how long participants fixated targets, there was a significant effect of condition on saccadic amplitude, *F*(40,2) = 7.73, *p* = .001, *η^2^* = 0.28. This was driven by a difference between the No AiTR condition (*M* = 6.1°, *SE* = 0.18°) and both the AiTR condition (*M* = 5.02°, *SE* = 0.22°), *t*(42) = 3.69, *p_tukey_* = 0.002, Cohen’s *d* = 1.37, and the Delayed AiTR condition (*M =* 5.20°, *SE* = 0.22°), *t*(42) = 3.04, *p_tukey_* = 0.01, Cohen’s *d* = 1.15. There was no significant difference in average saccadic amplitude between the AiTR condition and the Delayed AiTR condition.

### Situational awareness

#### Preregistered analyses

To analyze memory strength, a receiver operating characteristic (ROC) curve was calculated for each participant in each condition. This method avoids potential pitfalls of the more classic d-prime analysis approach by not assuming participants apply the same threshold criteria when deciding if a memory representation is sufficiently strong enough to report an item as having been seen (see Brady et al., (2023) for a detailed explanation). Having participants report on a continuous scale of old to new for each object allows memory to be tested at various threshold criteria. To do this, the continuous scale was discretized into 8, equally spaced, points. Then each point was in turn set as the criteria for classifying an item as ‘old’ or ‘new’ (e.g. all items given a score over this point were marked as reported ‘new’ and all items scored under this point were marked as ‘old’) and the true positive and false positive rates were calculated. This produces an ROC curve, where a straight line from the bottom left corner of the graph to the top right would indicate chance performance. To compare performance between conditions, the area under the curve (AUC) was calculated, where higher AUC indicates better memory. Averaging this value across participants within condition showed that numerically, the AiTR condition had a lower AUC (*M* = 0.68, *SE* = 0.02) than the No AiTR condition (*M* = 0.70, *SE* = 0.02) or the Delayed AiTR condition (*M* = 0.72, *SE* = 0.02). However a one-way ANOVA showed no significant difference between conditions, *F*(53,2) = 0.80, *p* = 0.45. None of the preregistered comparisons were significant.

#### Exploratory analyses

Previous work found that highly salient objects can make it into memory without being directly fixated (Bainbridge, Hall, & Baker, 2019). As such it could be that the higher salient items could enter into memory, regardless of the allocation of overt attention and thus mask the effect of condition. To explore this, the Graph Based Visual Saliency (GBVS) model (Harel, Koch, & Perona, 2006) was run on the scenes. For each item, in each scene a rectangular bounding box was drawn that contained edges of the object. The average saliency per pixel within the item’s bounding box was then calculated. This was then averaged across scenes, giving each item an average saliency score. A one-way ANOVA was then re-run, limiting the analysis to just reports made on items in the lowest salience quartile, as previous work shows higher saliency items may not require overt attention for encoding. The results followed a qualitatively similar pattern as the preregistered analysis with the average AUC of the AiTR condition (*M* = 0.72, *SE* = 0.06) being lower than both the No AiTR condition (*M* = 0.74, *SE* = 0.07) and the

Delayed AiTR condition (*M* = 0.78, *SE* = 0.08). However there was no significant difference across conditions, *F*(53,2) = 3.11, *p* = 0.05, *η^2^* = 0.10.

## Discussion

The human visual system is continuously engaged in processing large quantities of visual data. AiTR modifies this input by superimposing digital information onto physical objects in the real world. As a result, AiTR, and comparable systems in other fields (e.g. CAD systems), alter the distribution of bottom-up saliency across the visual field as well as up-weight highlighted items in their correspondence to task-goals. These alterations are by design as the system’s purpose is to encourage the human operator’s attention to task-relevant locations in the visual field, decreasing response times and increasing target detection rate. However, the design that elicits this enhanced effect for target detection may also compromise situational awareness and the likelihood of detecting critical targets missed by the system. As algorithm performance will never be completely accurate, whether due to adversarial attempts or incomplete information (e.g. a change in the human operators’ task goals), it is essential that while attentional deployment is influenced by AiTR highlights, this direction is not so complete as to abolish processing of unhighlighted information. The current work sought to use a combination of empirical findings of overt and covert scene processing as well as predictions of an attentional models (Wyble et al, 2020), to inform an implementation of AiTR highlights that would strike a balance between encouraging attentional resources to identified task-relevant locations while preserving natural processing of background information.

The first hypothesis of this experiment was that the presence of AiTR highlights would result in reduced attention towards background objects, resulting in a lower percentage of overall fixations to background information. This was supported by the results which showed a lower percentage of fixation made to background information in the AiTR condition compared to the no AITR condition. The second hypothesis predicted that due to this decreased attentional allocation, participants would exhibit lower situational awareness when AiTR highlights were present, as opposed to when they were absent. While the results qualitatively followed the predicted pattern of the second hypothesis, there was no significant difference in situational awareness, as measured by the surprise memory test, between the AiTR and No AiTR conditions. The third hypothesis posited that by delaying the presentation of AiTR highlights in relation to scene onset, there would be a greater allocation of attention to background objects and, as a result, participants would have better memory for those objects compared to participants who were presented AiTR highlights simultaneously with scene onset. Again, the attentional allocation portion of this hypothesis was supported by the results of this study as the Delayed AiTR condition showed an increase in the percentage of background fixations compared to the AiTR condition. However, this did not translate into a significant difference in memory performance for those background objects.

### Attentional allocation

Given what is known about the influence of both bottom-up signals and top-down goals on eye movements (e.g. Malcolm & Henderson, 2010; Orquin, Bagger, & Loose, 2013; Zelinsky et al., 2005) it is unsurprising that a lower percentage of fixations would be made on background information in the AiTR condition compared to the No AiTR condition, as AiTR highlights are both salient and task-relevant. However, what is novel about the current findings is that, in delaying the onset of AiTR highlights by only 250 ms, this decrease in background fixations was reversed. Typically, studies looking at temporal components of eye movements either delay when saccades are allowed (Talcott & Gaspelin, 2021) or analyze early versus late saccades to examine differences in processing (Mulckhuyse, Van de Stigchel, & Theewues, 2009). Here, the eyes were allowed to move freely in all conditions and the only difference in visual input between the AiTR condition and the Delayed AiTR condition was in the short period at the start of the trial before the highlights onset in the Delayed condition. This brief window (short enough that only 4% of all saccades made during the trial were executed prior to the onset of highlights) significantly altered where the eyes moved for the rest of the trial (where the average trial took 7.5s to complete). Models of attention that only consider bottom-up and top-down influences would not be able to account for these findings as, once the highlights were presented, the balance of those two factors across each scene was identical in the AiTR and Delayed AiTR groups. However, in a computational model of attention developed by Wyble and colleagues (2020) that simulates the temporal as well as spatial properties of attention, relative stimulus onset time is critical in determining whether it is attended. Namely, locations in the visual field can be thought of as being engaged in an attentional race. A location can win this race because it has a stronger priority (a combination of its bottom-up saliency and its top-down relevancy to task goals) or because it was given a sufficient temporal head start relative to higher priority areas. By demonstrating that future eye movements are influenced by a preceding difference in the visual input (the highlight delay), the current work provides evidence in support of this model prediction.

### Situational Awareness

Situational awareness, or one’s ability to attend and act on information in their environment, is obviously a critical construct for military purposes (and in other professions). However, it is a difficult one to operationalize. In the current work, situational awareness was operationalized using a surprise memory test of background objects. This method was chosen over more immediate probing of awareness (e.g. having participants respond to a peripheral cue during the search trials) as such probing introduces a secondary task and necessarily changes how participants perform search. Alternatively, other work has used more immediate surprise questions during a task (e.g. after asking participants to report the orientation of a stimulus on every preceding trial, the participant is given a surprise trial where they are asked to report the color of the stimulus instead as a means of testing what task-irrelevant information has been encoded (Swan, Collins, & Wyble, 2016)). However, this means that only a single trial of data is collected per participant. As the current work pulled from a special population (i.e. active-duty soldiers and veterans) and required in-person testing (due to the incorporation of eye tracking), the sample size was already smaller than originally planned. Therefore, the current method to operationalize situational awareness was used as it extracted the most information out of each participant. This was done not only by asking participants about all the items they saw, but also in having them report memory strength with a continuous scale. With a classic two forced-choice method that are analyzed using d-prime, differences between conditions can be driven either by true differences in the strength of memory representation or by a shift in reporting threshold (e.g. participants in one condition required less strong memory representations to report an object as “old” than participants in another condition) (Brady et al., 2023). The continuous memory scale allows for the comparison of memory strength across a variety of reporting thresholds, providing a better comparison of the true strength in memory across conditions. However, despite this method isolating the measure of interest, the expected difference between the No AiTR and AiTR conditions and between the AiTR and Delayed AiTR conditions, were not observed.

This lack of expected difference could be a power issue—the effect of condition on memory was too small to be observable in the available sample. However, another point that should be made is that this operationalization of situational awareness is highly conservative. Not being able to report having seen an item many minutes later, while performing an intervening search task, does not necessarily mean that one was not aware of the item when it was presented. Whether someone remembers something is a combination of factors including whether the information was encoded and whether it was successfully maintained during the retention period (Kim, 2019). Moreover, it is known that the brain is capable of discarding task-irrelevant information (Gaschler, Marewski, & Frensch, 2015). As such, the method used here may only have been able to analyze memory representations that were already sufficiently strong as to have survived until the surprise memory task. It could be that this unintentional thresholding is the reason that more variability was not captured and described by the manipulation of condition. The exploratory analysis, excluding items with high saliency, was an effort to eliminate items that, theoretically, could have been encoded and retained without being overtly attended. However as with the confirmatory analysis, while the results were in the direction predicted, this effect was not significant. Again, it could be that the longer retention period (between beginning the search task and beginning the memory task) already eliminated the weaker memory representations that may have been affected by highlight presentation. Another potential future direction could be to test memory not through an old/new choice but through a multiple alternative forced choice, as such paradigms may yield a more sensitive, unbiased measurement of memory strength (Brady et al., 2022).

### Conclusion

Visual search is a critical component for many professions. Whether it is a baggage screener searching x-ray images of luggage for prohibited items, a radiologist searching CT scans for abnormalities, or a soldier searching their surroundings for threats, failure to detect targets can have profound consequences. AiTR systems, and those similar, have the potential to limit search errors and speed response times to targets. However, to reach that potential, such systems will need to leverage the vast amount of research that has been done on how the human brain performs visual search. The current work sought to do this by testing a display presentation strategy informed by previous research and modeling, namely, delaying the onset of highlights. This delay proved effective in allowing participants to direct overt attention to unhighlighted regions. In tasks such as baggage screening or searching lung scans, where the onset time of information is more defined (e.g. x-ray images are presented sequentially on a screen), these results suggest that potentially presenting highlights after a brief delay could allow searchers to process images more wholistically before encouraging attentional resources to algorithmically-defined areas of interest. For professions where search is more of a continuous process, such as a soldier surveying the environment, a potential application of these results could be to have AiTR highlights presented intermittently, either by having the soldier themselves toggle the system on or off or have the system respond to commander input or environment sensors (e.g. turning off briefly as soldiers move from an indoor to outdoor). The unintended benefit of such a recommendation is that not having continuous AiTR displays would also help to address reported eye strain symptoms known to occur with current headsets (Hirzle, Fischbach, Karlbauer, & Jansen, 2022). Together, these results demonstrate how foundational research of the visual attention system can be used to develop AiTR systems that complement, rather than conflict with, the cognitive mechanisms of the human user.

## Declarations

### Consent for publication

Not applicable.

### Availability of data and material

Data generated and analyzed during the current study is available is at www.osf.io/3rka8 along with the preregistration, analysis code used and experimental paradigm code.

### Authors’ contributions

CCF was responsible for building the experimental paradigm, analysis and writing. AJ and JLV were responsible for subject recruitment and data collection. GBL was responsible for the original idea for the study. All authors contributed to, read and approved the final manuscript.

## Acknowledgements

The authors would like to acknowledge Alex Stauff who created with stimuli set.

